# Increased vaccine efficacy against tuberculosis with a recombinant BCG overexpressing the STING agonist cyclic di-AMP

**DOI:** 10.1101/2025.07.17.665241

**Authors:** Dhiraj K. Singh, Peter Um, Garima Arora, Xavier Alvarez, Vinay Shivanna, Edward Dick, Smriti Mehra, William R. Bishai

**Affiliations:** Southwest National Primate Research Center, Texas Biomedical Research Institute, San Antonio, TX, USA; Johns Hopkins School of Medicine, Center for TB Research, Baltimore, MD, USA

## Abstract

Tuberculosis (TB) remains the leading cause of death due to infection globally. Bacillus Calmette Guérin (BCG), a live attenuated bacterial strain, is the only available TB vaccine, but it has poor efficacy in preventing pulmonary TB in adults. There are advantages associated with the BCG platform however, including a remarkable safely profile, billions of administered doses and a public health ecosystem associated with its production, administration and care. A recombinant/modified BCG (rBCG/modBCG) anti-TB vaccine candidate would be able to leverage these advantages while improving on efficacy. BCG-STING is a recombinant BCG strain which overexpresses the mycobacterial diadenylate cyclase *disA* gene leading to the release of ∼15-fold greater levels of the endogenous small molecule STING agonist cyclic di-AMP. We evaluated vaccination with BCG-STING compared to its parental BCG-WT strain in rhesus macaques challenged with virulent *Mycobacterium tuberculosis* (Mtb). BCG-STING given intradermally was well-tolerated, and during life serial BAL samples showed that BCG-STING vaccinated animals had lower Mtb CFU counts than those receiving BCG-WT. At necropsy, BCG-STING vaccinated animals had significantly lower Mtb lung CFU counts, and a higher percentage of sterile lobes and granuloma lesions than the BCG-WT recipients. The enhanced protection observed in BCG-STING vaccinated animals was associated with significant elevations of antigen-specific CD4^+^ and CD8^+^ T cells. Modifying BCG to overexpress a small molecule recognized by the innate immune system significantly improves protection and cell-mediated immune responses against TB the non-human primate (NHP) model.

## INTRODUCTION

The BCG vaccine was introduced in 1921 for TB prevention, and WHO guidelines recommend that it be given to virtually all children at birth. In 1975 intravesical BCG was found to have potent antitumor activity against non-muscle invasive bladder cancer (NMIBC), and it is now a first-line therapy for NMIBC. Across these two important indications, BCG is believed to be the most widely administered vaccine in human history. Despite its widespread use, BCG has poor activity in preventing pulmonary TB in adults and intravesical BCG is unsuccessful in ∼30% of bladder cancer patients.^1,2^ Thus, there is an important unmet need for improved versions of BCG.

Many recombinant BCG strains have been developed for TB prevention, and typical strategies include overexpression of known Mtb antigens^3^, cytokines^4^ or adjuvants^5^, or partially restoring genes known to have been deleted in BCG.^6^ Despite the fact that many of these experimental recombinant BCG strains have shown improved protection against TB in small animal models, only two live attenuated mycobacterial vaccines have advanced to human clinical trials.^7,8^ Using a different approach to those described above, we are developing a modified BCG which overproduces a small immunostimulatory molecule: the stimulator of interferon genes (STING) agonist, cyclic di-AMP. Our recombinant BCG, therefore, should retain the classic properties of BCG itself but also have additional immunostimulatory properties by virtue of high-level STING agonist release.

Recent studies have identified a critical role for the stimulator of interferon genes (STING), an intracellular sensor, in mediating innate immune responses to cellular stress or pathogen infections.^9^ STING is a cytosolic receptor for both pathogen-associated molecular pattern (PAMP) molecules such as cyclic dinucleotides c-di-AMP or c-di-GMP produced by bacteria, and for endogenous mammalian danger signaling DAMP molecules such as 2’,3’ cyclic GMP-AMP (cGAMP). cGAMP is synthesized by cGAS (cyclic GMP-AMP synthase) in response to microbial or self-derived cytosolic double-stranded DNA.^9–11^ Activation of STING induces numerous interferon-stimulated genes including type I interferons (IFNα/β), and is associated with co-activation of the NF-ΚB and STAT6 transcription factors.^12^ Thus, endogenous and exogenous cyclic dinucleotides (CDNs) are strong TLR-independent mediators of innate host defenses. Based on the key role of STING signaling, pharmacological stimulation of the pathway using small molecule STING agonists may enhance anti-tumor and/or anti-viral immunity.^13,14^ CDN STING agonists also exhibit attractive vaccine adjuvant properties. They increase expression of MHC class II, co-stimulatory molecules (CD80/CD86) as well as activation (CD40) and adhesion (CD45) markers; in addition STING agonists have been shown to enhance antigen processing and presentation to T cells.^15–17^ Among microbial-derived CDNs, c-di-AMP has emerged as an efficient activator of macrophages and leads to robust Th1, Th17 and CD8 T cell responses.^18^

Previous studies from our group describe the construction of BCG-STING (also called BCG-*disA*-OE) and its efficacy for TB prevention in the guinea pig model^19^ and as a bladder cancer immunotherapy in mouse and rat models.^20^ In this report we describe the efficacy of BCG-STING for preventing TB disease progression in the Rhesus macaque non-human primate (NHP) model.^21,22^

## RESULTS

### Intradermal BCG-STING is as well tolerated as BCG-WT

Mycobacteria-free (negative TST, **Table S1**), specific pathogen-free, rhesus macaques ages 3 to 11 years were sourced from either the Tulane National Primate Research Center (TNPRC) or the California National Primate Research Center (CNPRC). On day 0, group 1 (5 males, 2 females) were left unvaccinated while the following intradermal vaccinations were given: BCG-Pasteur (BCG-WT) 1×10^6^ CFU-group 2 (3 males, 4 females), and BCG-STING (KanR gene-free version, known as OS-151) 1×10^6^ CFU - group 3 (8 males, 0 females), (**Fig. 1a**). Following vaccination, there were no skin ulcerations at the injection sites or signs of disease such as dyspnea, anorexia, pyrexia, body weight loss or changes in vital signs. Additionally, there were no changes pre-and post-vaccination in inflammatory or disease markers including mean body temperature (**Fig. S1a**), weight change (**Fig. S1b**), neutrophil/lymphocyte (N/L) ratio (**Fig. S1c**), CRP (**Fig. S1d**) or albumin/globulin (A/G) ratio (**Fig. S1e**).

**Figure 1.**
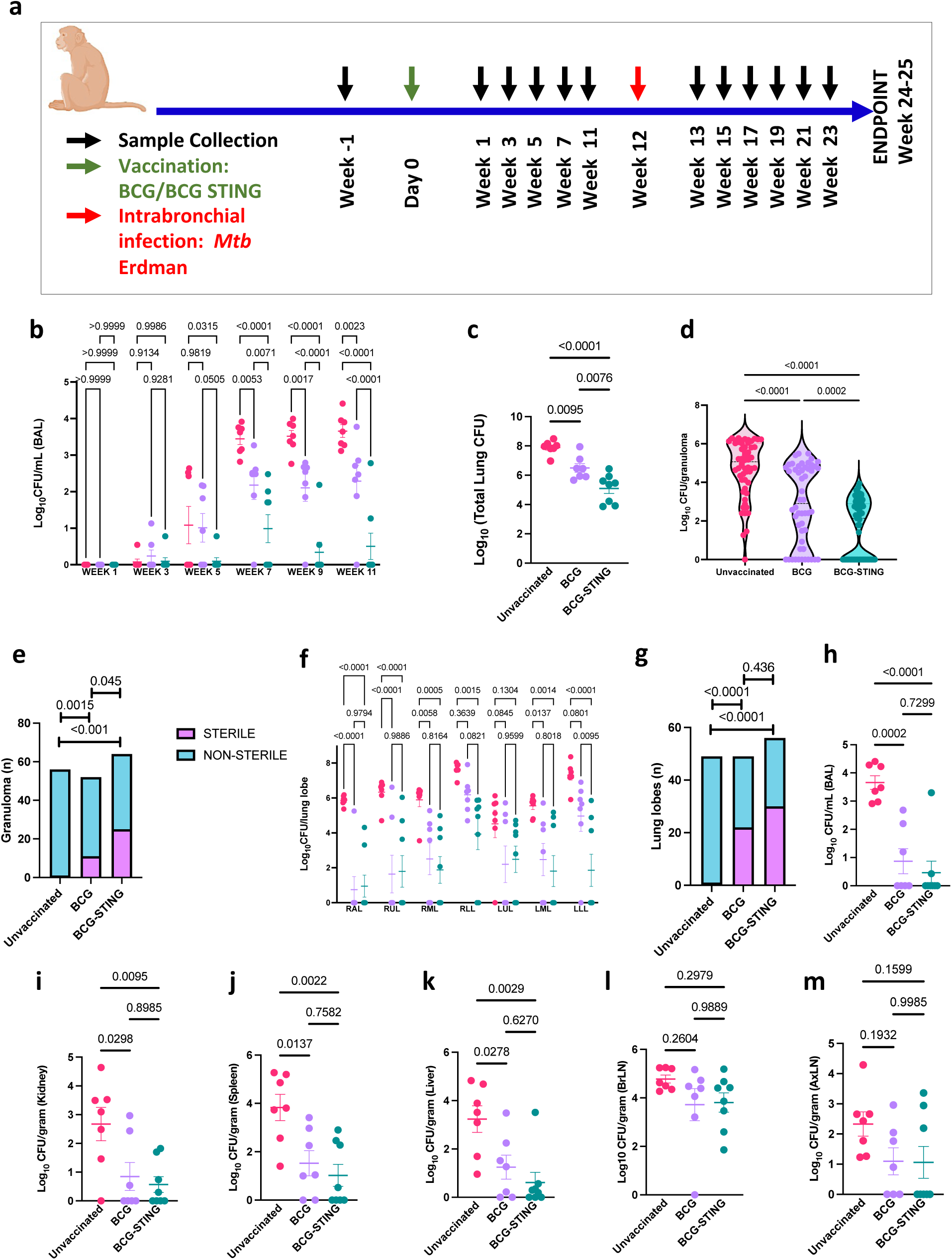
BCG-STING vaccination protects macaques from tuberculosis. (**a**) Experimental design and sampling schedule for testing vaccine efficacy in Unvaccinated (strawberry), BCG (lavender) and BCG-STING-vaccinated (teal) RMs challenged with *Mtb* Erdman. The schematics were created using BioRender. In unvaccinated (strawberry), ID BCG (lavender) and ID BCG-STING (teal) RMs post-Mtb infection, shown are Colony Forming Units (CFU) in (b) longitudinal BAL, (c) endpoint lung (total) and (d) granuloma. (e) Fisher’s exact test comparing sterile (purple) vs nonsterile (turquoise) granuloma. (f) CFU in endpoint lung lobes at endpoint and (g) Fisher’s exact test comparing sterile (purple) vs nonsterile (turquoise) lung lobes. CFU in endpoint (h) BAL, (i) Kidney, (j) Spleen, (k) Liver, (l) Bronchial and (m) axillary lymph nodes.

### Improved containment of Mtb burdens in the lungs and reduced extrapulmonary dissemination by BCG-STING-, compared with BCG-WT-vaccination

12 weeks post-vaccination, all NHPs were challenged with Mtb Erdman ∼10 CFU administered via intrabronchial instillation.^23^ TB disease progression was monitored by serial vital sign measurements, blood tests, and bronchoscopy with bronchoalveolar lavage (BAL) every two weeks (**Fig. 1a, S1f-j**), and PET-CT scans were performed at 4 and 12 weeks post-Mtb challenge. BAL sampling post-challenge revealed that beginning at week 5, BCG-STING vaccinated animals showed significantly lower BAL CFU counts than either BCG-WT or unvaccinated animals (**Fig. 1b**). Indeed, by week 7 the majority of BCG-STING-vaccinated NHPs showed sterile BAL washings (e.g. 5/8 or 62.5% at week 7) while no unvaccinated NHPs had sterile BAL samples at any time point from week 7 onwards and only 1 of 7 (14%) for BCG-WT showed sterility (weeks 7-11).

At necropsy 12-13 weeks post-challenge, we randomly collected 3-5 lung sections from each of the seven lung lobes, pooled them into a single paraffin-embedded block to yield seven lung blocks representing each lobe per animal for histological analyses. Adjacent lung pieces were collected and pooled per lobe along with 6-8 randomly dissected granulomas per animal for homogenization and quantitative CFU determination. Overall lung CFU counts for the BCG-STING NHPs were 5.1 log_10_ CFU units (**Fig. 1c**) which was significantly lower than for the BCG-WT (6.5) and unvaccinated groups (7.9). Similarly, the CFU counts per granuloma were significantly lower for BCG-STING animals (1.6 log_10_ CFU units) than for the BCG-WT (2.9) and unvaccinated (4.7) NHPs (**Fig. 1d**), and 39% of granulomas from BCG-STING animals were sterile as opposed to 21% from the BCG-WT (p 0.045) and only 1.78% unvaccinated groups (**Fig. 1e**). Across each lung lobe, CFU counts were lowest for BCG-STING-vaccinated and highest for unvaccinated animals (**Fig. 1f**), and both BCG-WT (44%) and BCG-STING vaccinated animals had significantly more sterile lung lobes (53%) than unvaccinated animals (2%) (**Fig. 1g**). We also considered the degree of Mtb dissemination to non-pulmonary organs. Both BCG-WT and BCG-STING animals has significantly lower kidney, spleen, and liver CFU counts than unvaccinated NHPs (**Fig. 1h-k**), and they both showed a trend towards reduced dissemination to bronchial and axillary lymph nodes compared to the unvaccinated group (p NS, **Fig. 1l,m**).

### Measurement of lung TB disease by PET/CT radiology and pathology

The week 4 post-challenge PET/CT scans showed that TB disease was limited to mediastinal lymph nodes in BCG-WT and BCG-STING vaccinated animals, while unvaccinated NHPs demonstrated more advanced infiltrative lung lesions (**Fig. 2a-c, Fig S2a-c**). By week 12 all animals had lung infiltrates, but unvaccinated NHPs had comparatively more lesions (**Fig. 2d-f, Fig S2d-f**). The granulomas detected by PET/CT showed significantly smaller volumes and abundance numbers in both of the vaccinated groups at week 4 post infection (**Fig. 2g)**. The SUVmax (Maximum Standardized Uptake Value), a measure of the metabolic activity of tissues, was also significantly lower in the vaccinated animals when compared to unvaccinated at this timepoint (**Fig. 2g)**.

**Figure 2.**
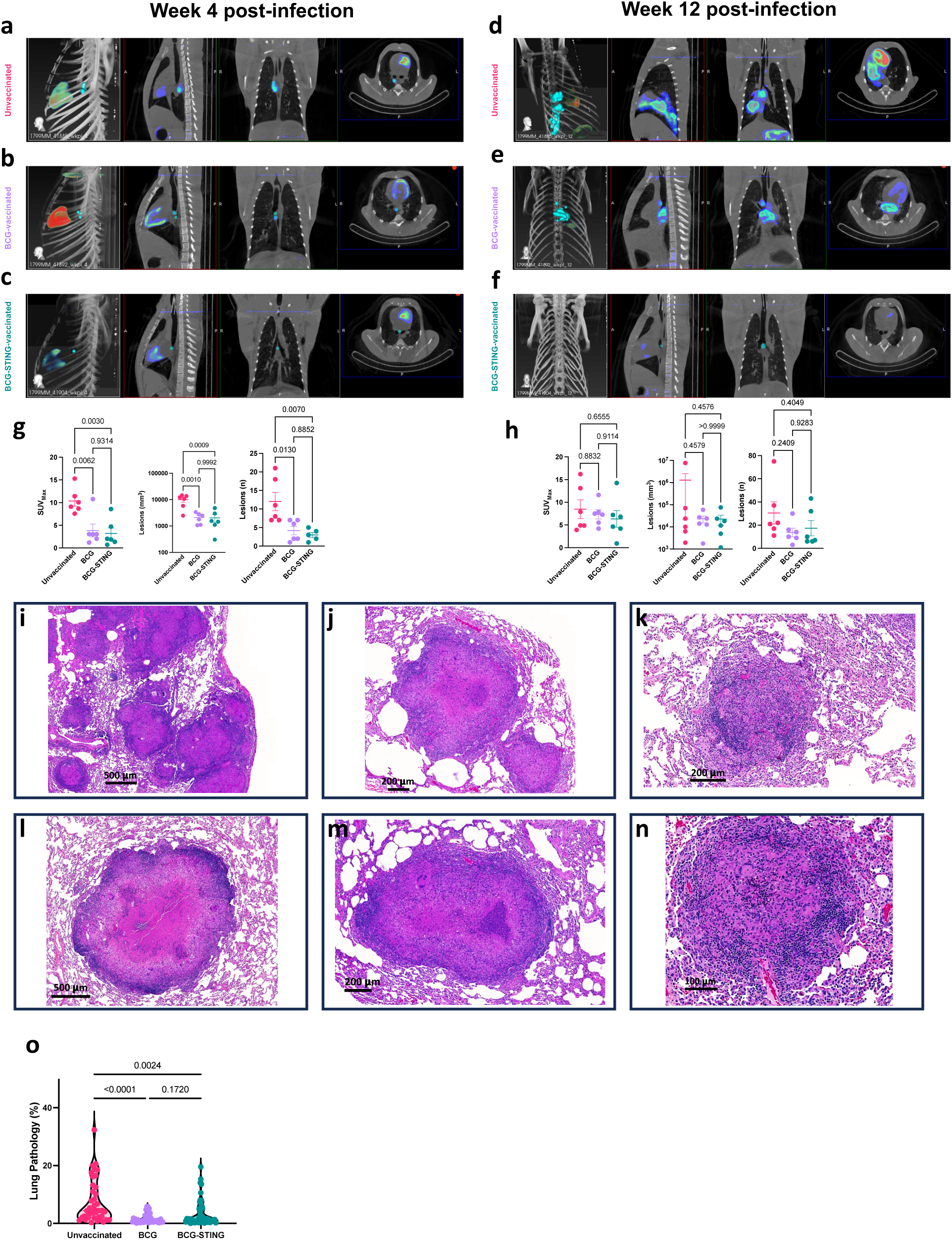
BCG-STING vaccination protects macaques from tuberculosis. Representative Positron Emission Tomography (PET) and Computed Tomography (CT) scans performed 4 weeks post-infection for (a) Unvaccinated, (b) BCG and (c) BCG-STING-vaccinated macaques showing (left to right) 3D reconstruction of the ROI volume representing the location of the lesions with transverse, vertical and longitudinal views. Representative PET-CT scans performed 12 weeks post-infection for (d) Unvaccinated, (e) BCG and (f) BCG-STING-vaccinated macaques showing (left to right) 3D reconstruction of the ROI volume representing the location of the lesions with transverse, vertical and longitudinal views. Shown are maximum standardized uptake value (SUV_Max_) (left), total volume of lesions (middle) and number of lesions quantified in PET-CT scans performed (g) 4 weeks post-infection and (h) 12 weeks post-infection in unvaccinated (strawberry), ID BCG (lavender) and ID BCG-STING(teal) vaccinated RMs. Representative H&E images showing histopathology and granulomatous pathology of unvaccinated (i,l), BCG-(j, m) and BCG-STING-vaccinated (k, l) RM lungs. Morphometric quantification of area containing lung pathology across multiple lobes of lung tissue in the three groups using HALO (Indica Labs).

Lung microscopic (**Fig. 2i-n**, **Fig. S3a-f**) and macroscopic (**Fig. S3g-l)** pathology showed a high degree of disease in unvaccinated animals (mean percent lung involvement 14.3%), and significantly lower disease scores were observed in BCG-STING and BCG-WT treated animals, although the BCG-STING and BCG-WT animals did not differ significantly from one another (**Fig. 2o**). The same was true in considering lung necrotic core area, and lung cellular inflammation area (**Fig. S3m,n**). Lung lymphocytic aggregates were significantly lower in BCG-WT-vaccinated NHP than either of the other two groups (**Fig. S3o**).

### Intradermal vaccination with BCG-STING elicits strong antigen-specific T cell responses in BAL

To evaluate the effects of our vaccines on cell-mediated immunity, we evaluated peripheral blood samples collected 5 weeks post-vaccination. PBMCs were exposed to *M. bovis* whole cell lysate (2 μg/mL) for 14 hours, and then flow cytometry was performed. Compared with BCG-WT, BCG-STING elicited significantly higher levels of Ag-specific-CD8 T cells expressing IFNγ, TNFα, granzyme-B, IL-2, and IL-17 and - CD4 T cells expressing IFNγ, TNFα, IL-2, and IL-17 (**Fig. 3a-j**). In contrast BCG-WT did not elicit improved Ag-specific responses compared to unvaccinated NHPs. Similar results were obtained using *M. tuberculosis* whole cell lysate (2μg/mL) (**Fig. S4a-j, S5**).

**Figure 3.**
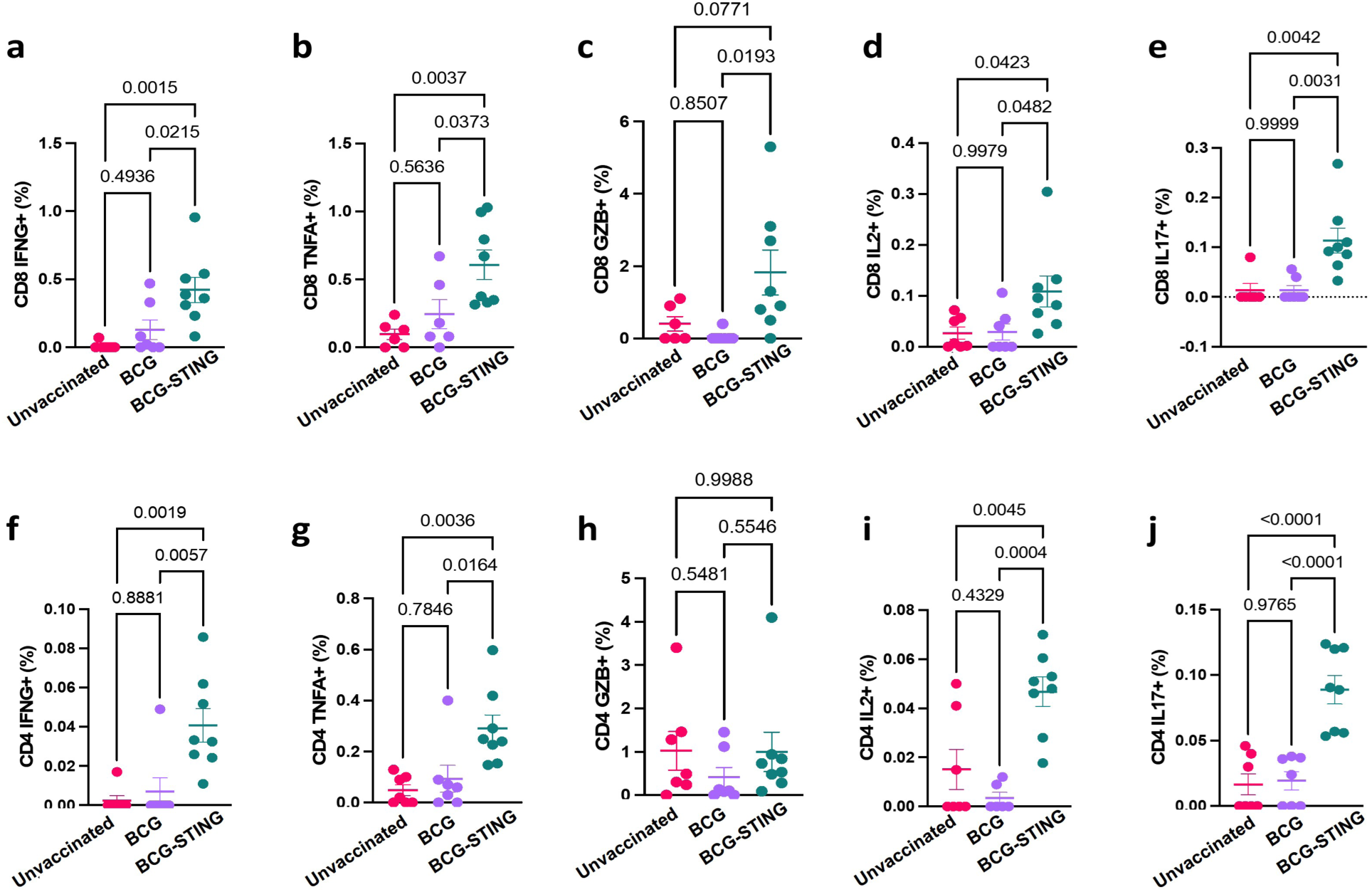
BCG-STING induces antigen specific T cell responses. Frequencies of CD8^+^ and CD4^+^ T cells expressing Interferon-γ (IFNG) (**a, f**), Tumor Necrosis Factor-α (TNFA) (**b, g**), Granzyme-B (GZB) (**c, h**), Interleukin-2 (IL2) (**d, i**) and Interleukin-17 (IL17) (**e, j**) in response to *M bovis* whole cell lysate in PBMCs collected 5 weeks post vaccination from unvaccinated (strawberry), with BCG-(lavender) or BCG-STING (teal) vaccinated macaques. Each dot represents an individual macaque.

## DISCUSSION

BCG-STING is a recombinant BCG strain which harbors a gene fusion between the strong constitutive hsp65 promoter (Phsp65) and the endogenous *disA* gene (encoding diadenylate cyclase) which generates the cyclic dinucleotide cyclic di-AMP^19,24^. Cyclic di-AMP is a mycobacterial second messenger, but is also detected as a PAMP by the STING-IRF3-TBK1 signaling pathway.^25^ While standard BCG strains deliver low levels of cyclic di-AMP to host cells (∼6 nM in macrophage cytosol), BCG-STING releases ∼15-fold higher levels. Indeed, BCG-STING has been shown to elicit higher pro-inflammatory cytokines, elevated levels of activated CD8 T cells, and an M1 myeloid cell shift in a STING-dependent manner.^20^ A previous vaccination studies in guinea pigs showed that intradermal BCG-STING confers a higher degree of containment of lung Mtb CFU.^19^

In the present study we found that despite engaging the STING pathway, known to stimulate potent NFΚB-mediated pro-inflammatory pathways, intradermal BCG-STING was well-tolerated in NHPs, and had an adverse effects profile indistinguishable from BCG-WT with both groups of animals showing normal vital signs and inflammatory markers post-vaccination. While earlier studies in immunocompetent (BALB/c) and in immunodeficient (SCID) revealed that BCG-STING was less pathogenic than BCG-WT, these safety observations in NHPs offer additional and important reassurance that BCG-STING does not elicit systemic hyperinflammation or cytokine storm.

Our vaccine efficacy studies showed that intradermal BCG-STING vaccination reduces detectable Mtb CFU counts in serial BAL samples to significantly lower levels than observed with BCG-WT. Moreover, at necropsy BCG-STING vaccinated animals had significantly lower total lung CFU counts, lower CFU counts per granuloma, and a higher abundance of sterile lung granulomas than in BCG-WT recipients. Beginning at 5 weeks post vaccination, BCG-STING vaccinated NHPs showed significantly elevated antigen-specific CD4 and CD8 T cell responses detected in PBMCs.

Previous studies of experimental TB vaccines in NHPs have often used intradermal BCG-WT as a control. These studies typically find that BCG-WT elicits a ∼1.0 log_10_ CFU reduction in total lung CFU compared to unvaccinated controls,^26–29^ although some studies observed more modest reductions in the range of 0.3 to 0.7 log_10_ CFU reductions.^30,31^ Our wild-type BCG strain (BCG-Pasteur) delivered intradermally conferred protection slightly higher than most historical comparators with a 1.4 x log_10_ CFU reduction compared to the unvaccinated animals (6.5 vs. 7.9 log_10_ CFU) while BCG-STING conferred an additional 1.4 x log_10_ reduction in total lung CFU beyond that of BCG-WT (5.1 vs. 6.5 log_10_ CFU). This relatively high degree of protection by our BCG-WT is a factor that would tend to reduce comparative efficacy of BCG-STING. Another important note is that we inadvertently assigned all male animals to the BCG-STING group (8 of 8), while the BCG-WT (3 M, 4 F) and unvaccinated (5M, 2 F) groups were more evenly split. Human studies have shown that females are more resistant to TB disease than males (M:F incidence ratio of active TB is 1.7:1)^32^ and are better protected against TB following BCG vaccination.^33,34^ This is a second factor which would tend to reduce the apparent efficacy of BCG-STING. An additional limitation of our study is that neither BCG-STING nor BCG-WT showed improved lung pathology at necropsy or PET-CT scores at the 12 weeks post-challenge, and there is a growing recognition that lung pathology may be an important contributor to post-TB lung disease.^35^

Highlighting the benefits of the BCG platform, dozens of recombinant BCG strains have been developed over the years, and many have shown improved efficacy over BCG-WT in TB prevention in small animal models.^8^ Using the prime-boost method with BCG as prime inoculation, several experimental TB vaccines now in clinical trials have been shown to confer superior reductions of lung CFU protection over that of BCG alone.^30,31^ It has also been shown that when BCG-WT or experimental vaccines are administered to NHPs by alternative routes such as endobronchially,^26^ by aerosol,^36^ or intravenously,^27,37^ lung CFUs and disease parameters can be significantly reduced to greater degrees than with intradermal BCG-WT. Despite these advances, it has proven difficult to show superiority to intradermally administered BCG in the NHP models. Thus, our study identifies BCG-STING as a promising experimental vaccine for TB prevention, the only one in the rBCG/modBCG space which may offer efficacy advantages over BCG-WT in the NHP TB model. Further preclinical and clinical development of this candidate vaccine may fill an important void in the current TB vaccine pipeline.

## METHODS

### NHP study design and infections

All procedures adhered to NIH guidelines and received approval from the Institutional Animal Care and Use Committees (IACUC) of Texas Biomedical Research Institute. Mycobacteria-naive rhesus macaques (RMs) obtained from the Tulane National Primate Research Center or California National Primate Research Center were used in this study protocol (**Table S1**). Specifically, the animals were either unvaccinated (n=7) or vaccinated intradermally with 1 x 10^6^ Colony Forming Units (CFUs) of BCG (n=7) or BCG-STING (n=8) (Figure 1a). 12-weeks post-vaccination, macaques were challenged with ∼10 CFU *Mtb* Erdman delivered to the lower right lung lobe via intrabronchial instillation^23^ using Ambu® aScope™ 4 Broncho attached to Ambu® aView™ 2 Advance unit. Infection was assessed through tuberculin skin test (TST), while TB progression was monitored by longitudinal measurements of weight, temperature, and C-reactive protein (CRP) and bronchoalveolar lavage (BAL) CFUs, and PET-CT (n=6 per group) as described (Figure 1a)^22,38–42^. Dissemination was evaluated during necropsy by culturing bronchial lymph node, spleen, liver, and kidney tissues to measure CFUs. Demographic information including age, gender, etc., and study specific information of macaques are provided (**Table S1)**. Animals were euthanized at 12-13 weeks post-challenge.

### Sampling

TST was performed before enrollment and at weeks 5 and 7 post-challenge, as described^22,42,43^. PET-CT scans were performed at 4 and 12 weeks post-infection as described^44^ and scored in a blinded manner. BAL and peripheral blood mononuclear cells (PBMC) samples were obtained one week before either vaccination or *Mtb* infection and subsequently every two weeks, as described^22,45^. BAL cells were used for determining bacterial burden and PBMCs were used for cellular analysis through flow cytometry, as described^22,42,45^. Blood samples were collected one week prior to vaccination or *Mtb* infection and thereafter on a weekly basis, for measuring complete blood count, serum chemistry, including serum C-reactive protein (CRP), and for flow cytometry, employing the flow panels specified in **Table S2**^40,42,45,46^.

### Tissue bacterial burden and pathology

Tissues were collected and processed as described^22,42^. CFUs were determined per gram of tissue and per mL of BAL fluid. Lung pathology at necropsy was assessed by a board-certified veterinary pathologist in a blinded manner, utilizing zinc-formalin-fixed paraffin-embedded (FFPE) tissues representing all lung lobes using previously described methods^22,42^.

### Immune analysis

Antigen specific T cell populations were quantified and characterized in PBMCs using flow cytometry, following established protocols^42,44,47–50^. Briefly, T cell populations and their antigen-specific functionality were assessed by stimulating PBMCs with *M. bovis* whole cell lysate (2μg/mL) (BEI Resources) or *Mtb* whole cell lysate (BEI Resources) for 14 hours, with addition of brefeldin A (Biolegend, USA) in the last 6 hours and analyzed using flow cytometry (**Table S2-3**, **Fig S5**), as detailed in prior publications^22,40,45,51^.

## Data Availability

Source data are provided with this paper. All data supporting the findings of this study are available within this manuscript and its Supplementary Information. Any additional data can be requested from the corresponding authors upon reasonable request.

## Supporting information

All supplemental tables

## Acknowledgments

This research was supported by NIH grant R01AI155346 to WB/SM and institutional NIH grants– OD011133, OD010442, OD032443, OD028732, OD028653 and OD028732. The content is solely the responsibility of the authors and does not necessarily represent the official views of the National Institutes of Health. *Mtb* Erdman, *M. bovis* whole cell lysate and *Mtb* whole cell lysate was obtained through BEI Resources, NIAID, NIH. BCG-Pasteur was provided by Dr. Frank Collins (FDA). We would like to thank Dr. Deepak Kaushal for providing critical insights through the manuscript.

## Author Contributions Statement

WB, SM, DKS, and PU designed the NHP study; DKS oversaw the NHP experiments, conducted research and data analysis with support from PU; GA; XA, VS, ED, Jr, contributed to research and/or analyses; WB, SM, DKS, and PU wrote the paper with help from all co-authors. All authors contributed to, edited and approved the manuscript. WB and SM provided funding.

## Supplemental Figures

**Figure S1.**
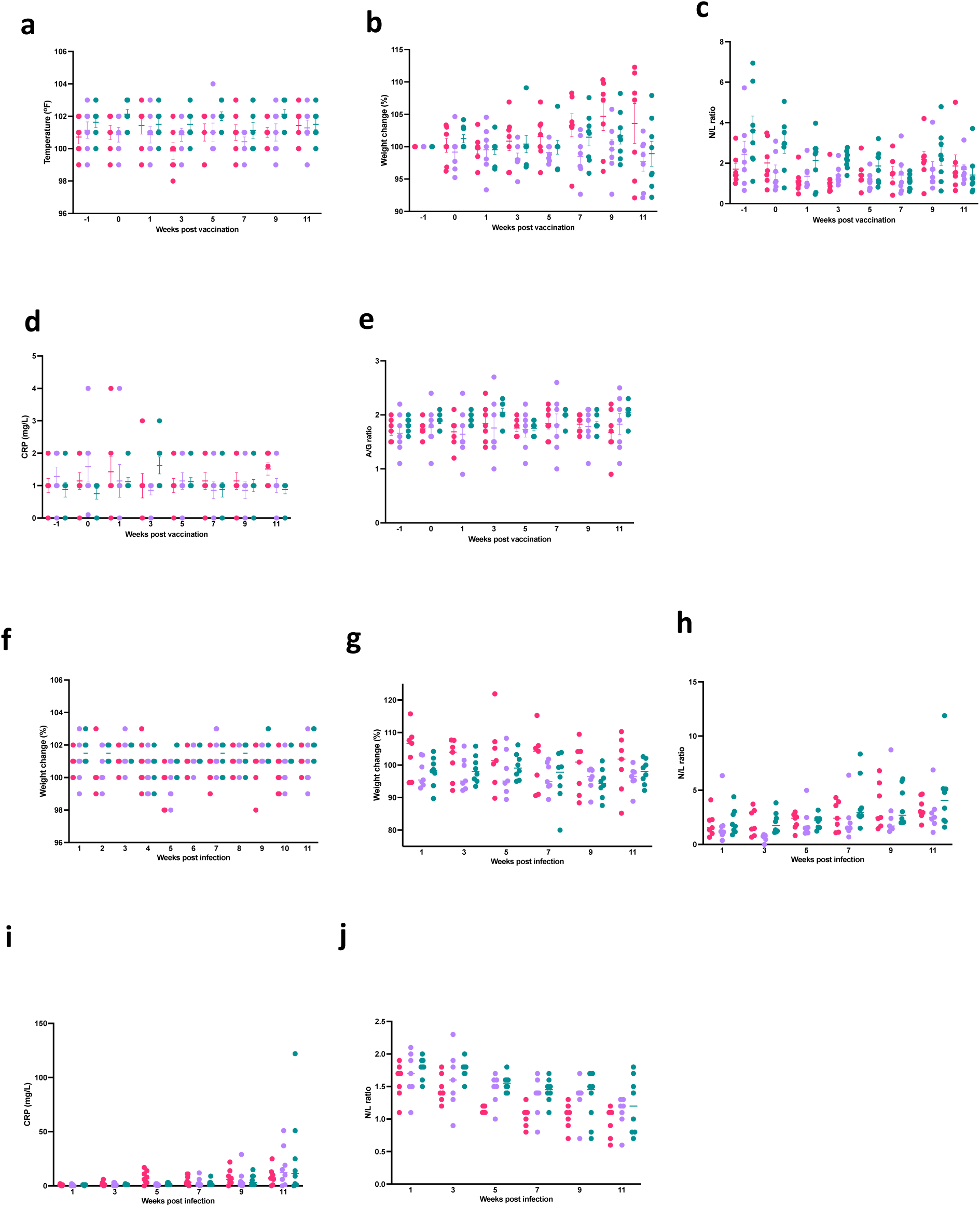
Clinical parameters post vaccination and infection. Shown are temperature (^0^F) (a), change in weight (%) relative to pre-vaccination (b), blood Neutrophil (N)/Lymphocyte (L) ratios (c), serum C-reactive protein (mg/L) (d) and Albumin/Globulin (A/G) ratios (e) post-vaccination in unvaccinated (strawberry), with BCG-(**lavender**) or BCG-STING (**teal**) vaccinated macaques. Shown are temperature (^0^F) (f), change in weight (%) relative to pre-vaccination (g), blood Neutrophil (N)/Lymphocyte (L) ratios (h), serum C-reactive protein (mg/L) (i) and Albumin/Globulin (A/G) ratios (j) post-infection in unvaccinated (strawberry), with BCG-(lavender) or BCG-STING (teal) vaccinated macaques.

**Figure S2.**
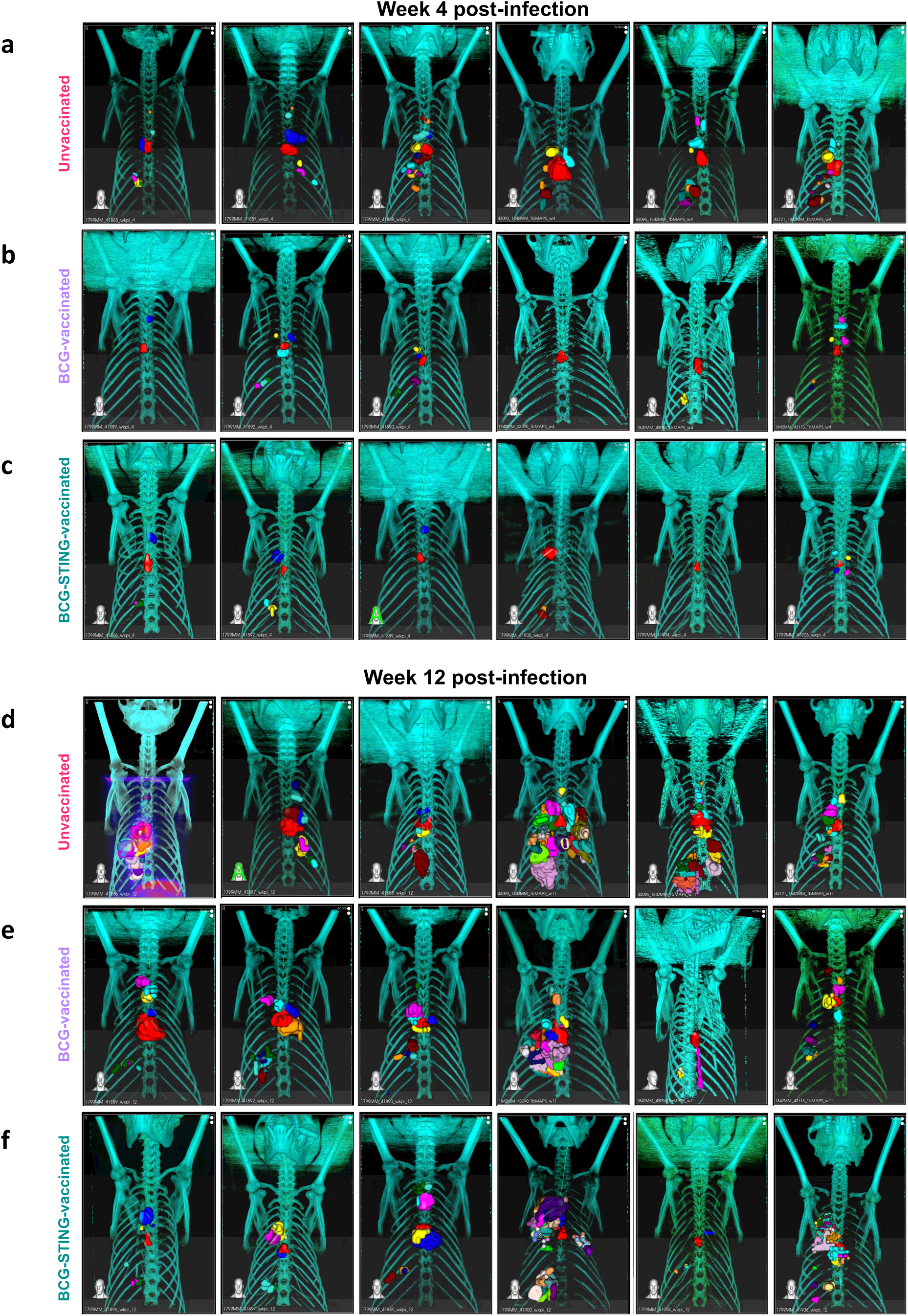
Representative Positron Emission Tomography (PET) and Computed Tomography (CT) scans showing 3D reconstruction of the ROI volume representing the location of the lesions performed 4 weeks post-infection for (a) Unvaccinated, (b) BCG and (c) BCG-STING-vaccinated macaques. Representative Positron Emission Tomography (PET) and Computed Tomography (CT) scans showing 3D reconstruction of the ROI volume representing the location of the lesions performed 12 weeks post-infection for (d) Unvaccinated, (e) BCG and (f) BCG-STING-vaccinated macaques.

**Figure S3.**
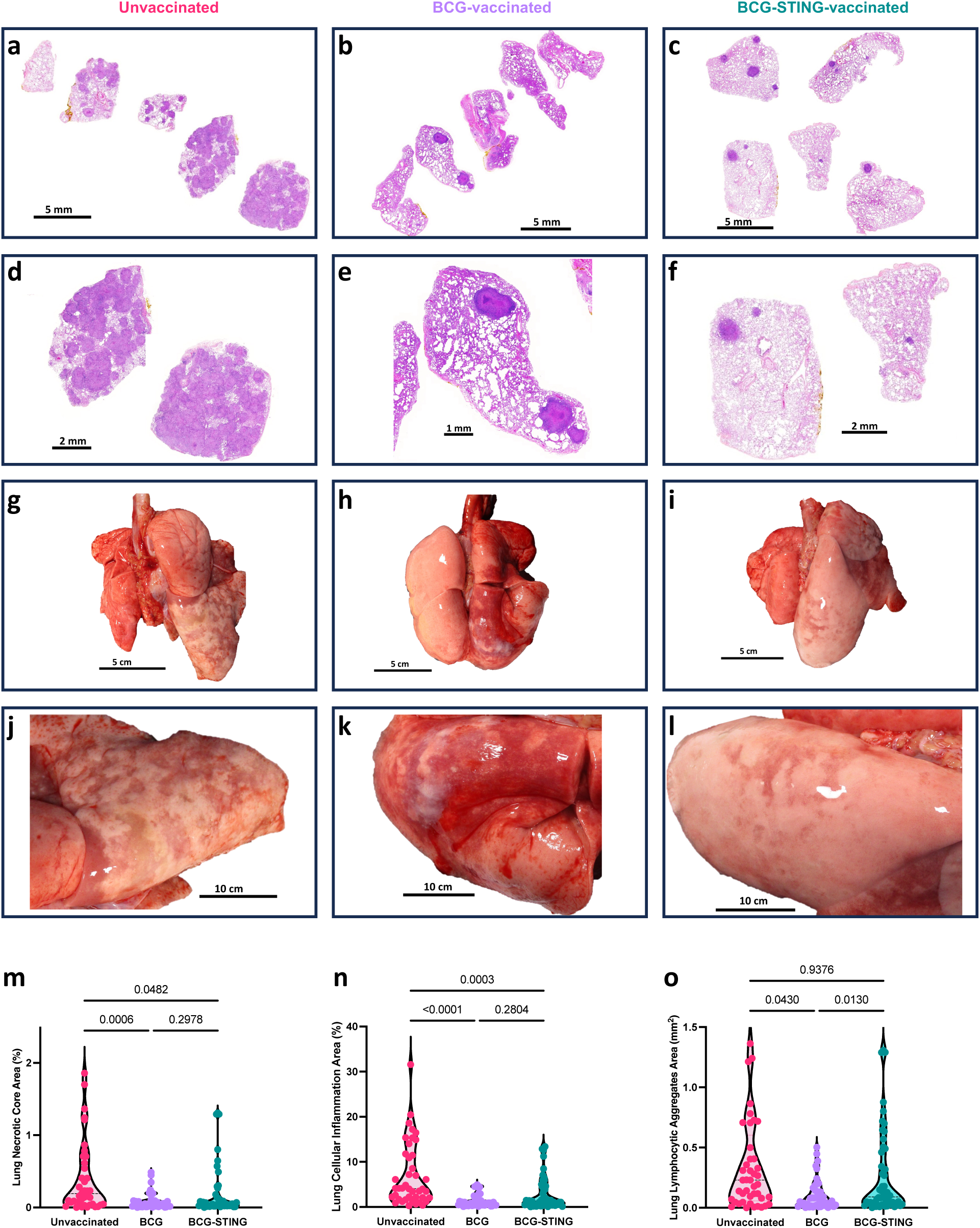
Representative H&E images showing lung histopathology (a-f) and gross pathology for lung (g-l) at different magnifications of unvaccinated (a, d, g, j), BCG-(b, e, h, k) and BCG-STING-vaccinated (c,f,i,l). Morphometric quantification of area containing necrotic lesion (m), cellular inflammation (n) and lymphocytic aggregates (o) across multiple lobes of lung tissue in the three groups using HALO (Indica Labs).

**Figure S4.**
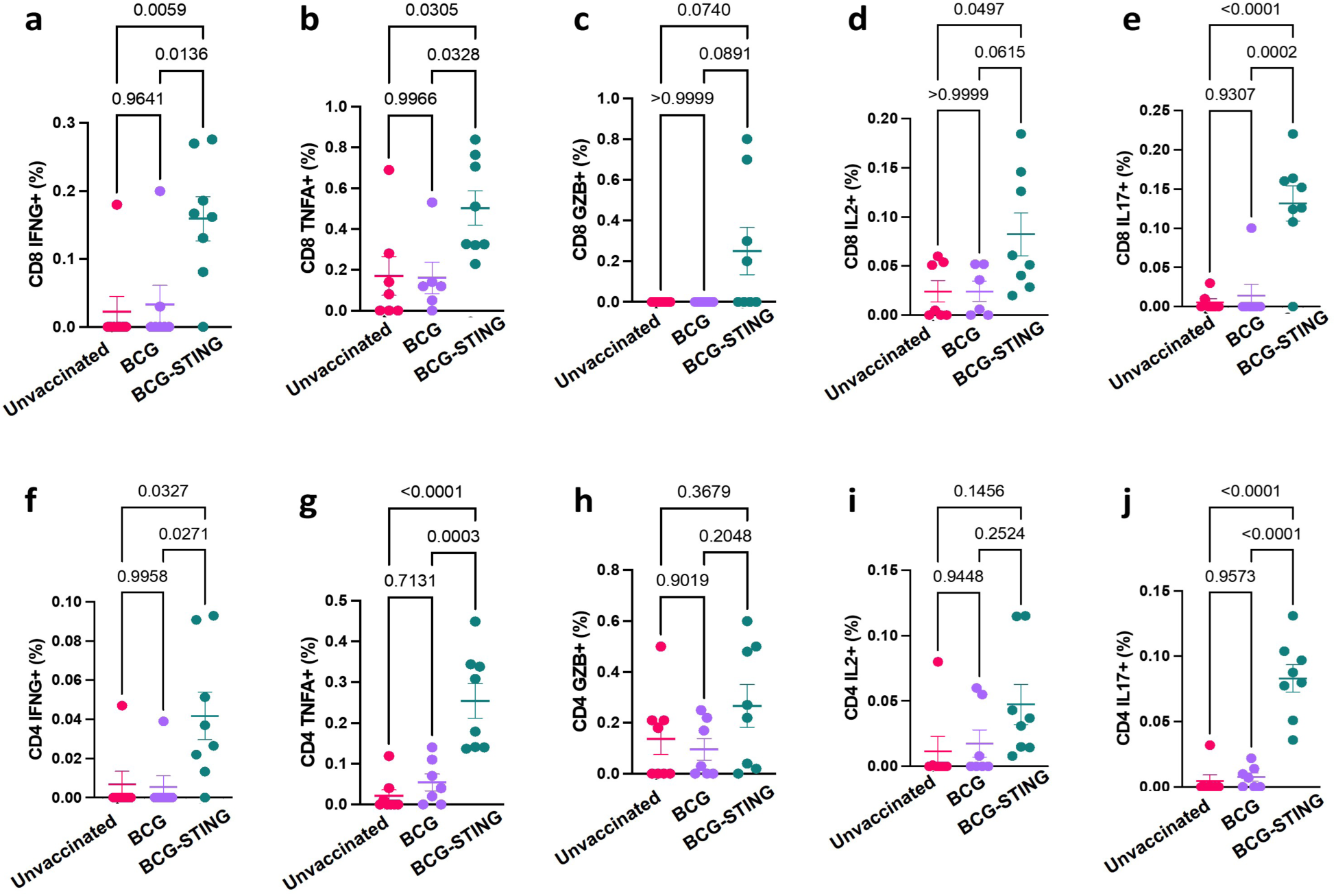
BCG-STING induces antigen specific T cell responses. Frequencies of CD8^+^ and CD4^+^ T cells expressing Interferon-γ (IFNG) (**a, f**), Tumor Necrosis Factor-α (TNFA) (**b, g**), Granzyme-B (GZB) (**c, h**), Interleukin-2 (IL2) (**d, i**) and Interleukin-17 (IL17) (**e, j**) in response to *Mtb* whole cell lysate in PBMCs collected 5 weeks post vaccination from unvaccinated (strawberry), with BCG-(**lavender**) or BCG-STING (**teal**) vaccinated macaques. Each dot represents an individual macaque.

**Figure S5.**
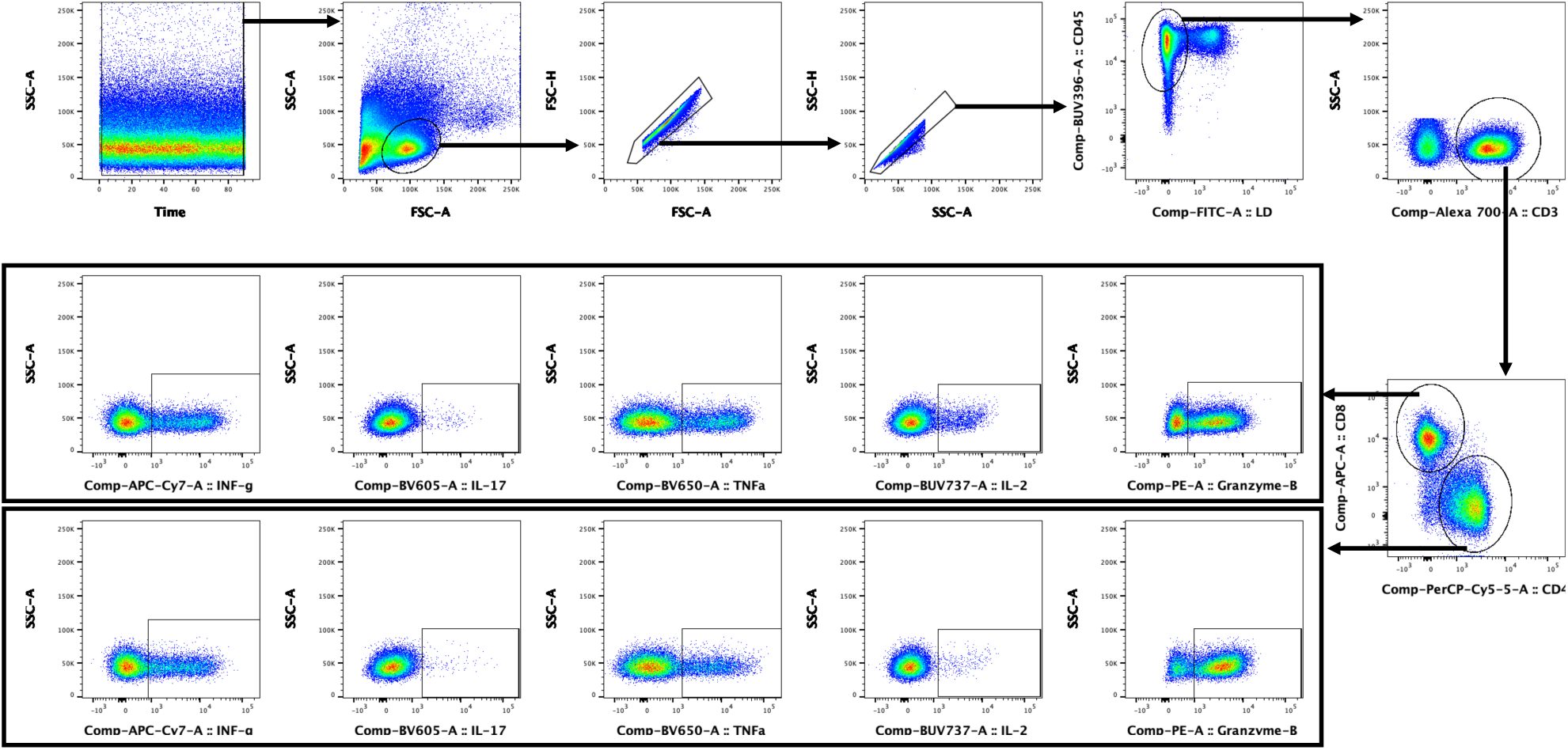
**Gating strategy for antigen specific T cell responses.**

