## Supplementary material for "Increased vaccine efficacy against tuberculosis with a recombinant BCG overexpressing the STING agonist cyclic di-AMP": All supplemental tables

Supplementary Table 1. Demographics for individual animals used in the study.

| Subject | Species | Gender | Age | Weight (kg) | Group | TST Status pre-vaccination | TST status (5 weeks post *Mtb*-infection) | Vaccination | Infection | Source |
|---|---|---|---|---|---|---|---|---|---|---|
| 41873 | *Macaca mulatta* | M | 5 years, 9 months | 10.00 | Unvaccinated | Negative | Positive | NA | Mtb Erdman | TNPRC |
| 41885 | *Macaca mulatta* | M | 4 years, 9 months | 6.00 | Unvaccinated | Negative | Positive | NA | Mtb Erdman | TNPRC |
| 41887 | *Macaca mulatta* | M | 4 years, 9 months | 6.60 | Unvaccinated | Negative | Positive | NA | Mtb Erdman | TNPRC |
| 41888 | *Macaca mulatta* | M | 4 years, 9 months | 5.80 | Unvaccinated | Negative | Positive | NA | Mtb Erdman | TNPRC |
| 43093 | *Macaca mulatta* | F | 5 years, 10 months | 6.03 | Unvaccinated | Negative | Positive | NA | Mtb Erdman | CNPRC |
| 43096 | *Macaca mulatta* | F | 4 years, 10 months | 5.83 | Unvaccinated | Negative | Positive | NA | Mtb Erdman | CNPRC |
| 43121 | *Macaca mulatta* | M | 3 years, 8 months | 5.71 | Unvaccinated | Negative | Positive | NA | Mtb Erdman | CNPRC |
| 41889 | *Macaca mulatta* | M | 4 years, 9 months | 8.60 | BCG | Negative | Positive | BCG | Mtb Erdman | TNPRC |
| 41890 | *Macaca mulatta* | F | 4 years, 9 months | 7.20 | BCG | Negative | Positive | BCG | Mtb Erdman | TNPRC |
| 41892 | *Macaca mulatta* | M | 4 years, 9 months | 7.40 | BCG | Negative | Positive | BCG | Mtb Erdman | TNPRC |
| 43090 | *Macaca mulatta* | F | 11 years, 10 months | 8.13 | BCG | Negative | Positive | BCG | Mtb Erdman | CNPRC |
| 43091 | *Macaca mulatta* | F | 10 years, 10 months | 10.56 | BCG | Negative | Positive | BCG | Mtb Erdman | CNPRC |
| 43094 | *Macaca mulatta* | F | 5 years, 8 months | 4.96 | BCG | Negative | Positive | BCG | Mtb Erdman | CNPRC |
| 43113 | *Macaca mulatta* | M | 4 years, 8 months | 7.36 | BCG | Negative | Positive | BCG | Mtb Erdman | CNPRC |
| 41893 | *Macaca mulatta* | M | 4 years, 8 months | 7.20 | BCG-STING | Negative | Positive | BCG-STING | Mtb Erdman | TNPRC |
| 41896 | *Macaca mulatta* | M | 4 years, 9 months | 6.00 | BCG-STING | Negative | Positive | BCG-STING | Mtb Erdman | TNPRC |
| 41897 | *Macaca mulatta* | M | 4 years, 8 months | 5.60 | BCG-STING | Negative | Positive | BCG-STING | Mtb Erdman | TNPRC |
| 41899 | *Macaca mulatta* | M | 4 years, 8 months | 6.20 | BCG-STING | Negative | Positive | BCG-STING | Mtb Erdman | TNPRC |
| 41900 | *Macaca mulatta* | M | 4 years, 8 months | 6.40 | BCG-STING | Negative | Positive | BCG-STING | Mtb Erdman | TNPRC |
| 41904 | *Macaca mulatta* | M | 4 years, 7 months | 6.60 | BCG-STING | Negative | Positive | BCG-STING | Mtb Erdman | TNPRC |
| 41905 | *Macaca mulatta* | M | 4 years, 6 months | 7.20 | BCG-STING | Negative | Positive | BCG-STING | Mtb Erdman | TNPRC |
| 41906 | *Macaca mulatta* | M | 4 years, 5 months | 5.80 | BCG-STING | Negative | Positive | BCG-STING | Mtb Erdman | TNPRC |

| Supplementary Table 2. Flow cytometry panel for measuring antigen specific T cell responses. | | | |
|---|---|---|---|
| S. No. | Marker | Flourochrome | Clone |
| 1 | CD45 | BUV395 | D058-1283 |
| 2 | IL-2 | BUV737 | MQ1-17H12 |
| 3 | CD95 | BV421 | DX2 |
| 4 | IL-17 | BV605 | BL168 |
| 5 | TNF-a | BV650 | Mab11 |
| 6 | Live/Dead | FITC | |
| 7 | CD4 | PerCP-Cy5.5 | L200 |
| 8 | Granzyme B | PE | GB11 |
| 9 | CD28 | PE-Cy7 | CD28.2 |
| 10 | CD8 | APC | RPA-T8 |
| 11 | CD3 | AL700 | SP34-2 |
| 12 | IFN-g | APC-CY7 | B27 |

| | | | S |
|---|---|---|---|
| **Antibody** | **Supplier** | **Clone** | **Cat Number** |
| CD45 (BUV395) | BD Bioscience | D058-1283 | 564099 |
| IL-2 (BUV737) | BD Bioscience | MQ1-17H12 | 612836 |
| CD95 (BV 421) | BD Bioscience | DX2 | 562616 |
| IL-17 (BV605) | Biolegend | BL168 | 512326 |
| TNF-alpha (BV650) | Biolegend | MAb11 | 502938 |
| CD4 (PCP-Cy5.5) | BD Bioscience | L200 | 552838 |
| GrB (PE) | BD Bioscience | GB11 | 561142 |
| CD28 (PE-Cy7) | BD Bioscience | CD28.2 | 560684 |
| CD8 (APC) | Biolegend | RPA-T8 | 301049 |
| CD3 (AL700) | BD Bioscience | SP34-2 | 557917 |
| IFN-G (APC-Cy7) | Biolegend | B27 | 506524 |

Supplementary Table 3. List of antibodies.

| Validation Statement: Reactivity |
|---|
| Rhesus, Cynomolgus, Baboon (QC Testing) |
| Human (QC Testing) Rhesus, Cynomolgus, Baboon (Tested in Development) |
| Human (QC Testing) Rhesus, Cynomolgus, Baboon (Tested in Development) |
| Human, Rhesus (Validated in lab, Reported) |
| Human, Cat (Feline)11 Cross-Reactivity: Chimpanzee, Baboon, Cynomolgus, Rhesus, Pigtailed Macaque, Sooty Mangabey, Swine (Pig, Porcine) |
| Rhesus, Cynomolgus, Baboon (QC Testing) Human (Tested in Development) |
| Human (QC Testing), Rhesus (NHP Reagent Resource) |
| Human (QC Testing), Rhesus (NHP Reagent Resource) |
| Chimpanzee, Baboon, Cynomolgus, Rhesus, Pigtailed Macaque, Sooty Mangabey |
| Rhesus, Cynomolgus, Baboon (QC Testing) Human (Tested in Development) |
| Chimpanzee, Baboon, Cynomolgus, Rhesus, Pigtailed Macaque, African Green, Sooty Mangabey |

| |
|---|
| Dilution |
| As recommended by Manufacturer |
| As recommended by Manufacturer |
| As recommended by Manufacturer |
| As recommended by Manufacturer |
| As recommended by Manufacturer |
| As recommended by Manufacturer |
| As recommended by Manufacturer |
| As recommended by Manufacturer |
| As recommended by Manufacturer |
| As recommended by Manufacturer |
| As recommended by Manufacturer |
